# The human parasite, *Toxoplasma gondii,* is paralyzed without two components of the apical polar ring

**DOI:** 10.64898/2026.03.07.707822

**Authors:** Jonathan Munera Lopez, Luisa F. Arias Padilla, Isadonna F. Tengganu, Yan Hao, Ying Zhang, Laurence Florens, John M. Murray, Ke Hu

**Author notes:** equal contribution (Jonathan Munera Lopez, Luisa F. Arias Padilla).

## Abstract

The phylum Apicomplexa contains ∼ 6000 known species of unicellular eukaryotic parasites. A unifying feature among the apicomplexans is the apical complex, which varies in complexity in different lineages, but always contains an annulus (a.k.a. the apical polar ring) into which the minus ends of an array of cortical microtubules are embedded. In *Toxoplasma gondii,* the apical complex also includes the conoid, which contains several signaling and structural proteins critical for parasite motility. The conoid extends and retracts through the apical polar ring in a calcium-dependent manner. Here we report the identification of several new apical polar ring components, including APR9, which is highly conserved among the apicomplexans and their free-living relative *Chromera velia*. The loss of APR9 alone has only a moderate impact on the parasite lytic cycle. However, the knockout of both APR9 and KinesinA (another apical polar ring component) paralyzes parasite and drastically impairs invasion, egress and the lytic cycle. The double-knockout displays multiple subcellular abnormalities, including the formation of an apical actin concentration, impaired conoid extension, and significantly reduced secretion of a major adhesin (MIC2) upon stimulation with a calcium ionophore. These findings reveal that the apical polar ring plays a critical role in parasite motility and contributes to multiple subcellular processes.

## INTRODUCTION

The apicomplexans are eukaryotic parasites specialized in infecting other eukaryotic cells. These parasitic protists infect many different types of metazoans [1]. Notable human pathogens among the apicomplexans include *Cryptosporidium*, *Toxoplasma*, and *Plasmodium* [1-4]. While the host range differs greatly among the apicomplexans, they are unified by key genetic and cellular architectures, the most iconic of which is the apical complex (**Fig 1A**), the eponym of the phylum.

**Fig. 1.**
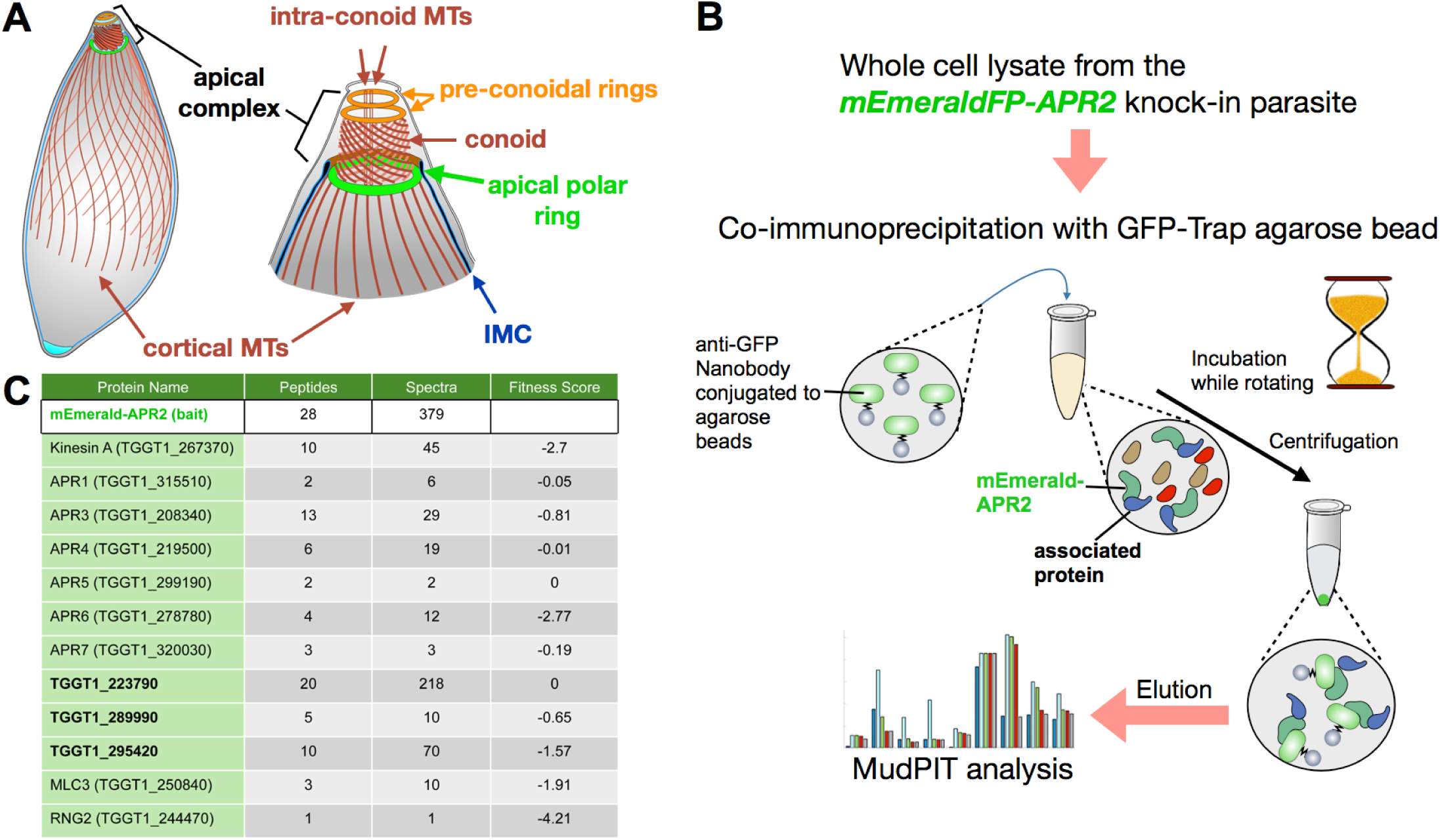
Identification of candidate apical polar ring proteins by immunoprecipitation and mass spectrometry analysis. **A.** Diagrams (modified from [26]) illustrating the organization of the tubulin-containing cytoskeleton and associated structures in *T. gondii*. IMC: Inner membrane complex. **B.** Diagram of the workflow for enrichment and identification of candidate apical polar ring proteins from the mEmeraldFP-*APR2* knock-in parasite line. mEmeraldFP-APR2 was used as the bait in immunoprecipitation (IP) with the high-affinity llama anti-FP antibody conjugated to agarose beads ("GFP-trap"). **C.** Table of peptide counts, unique spectral counts, and fitness scores for known apical polar ring proteins and for TgGT1_295420, 223790, 289990, identified by MudPIT in the IP. No spectra were detected for these proteins in the negative control (untagged WT parental parasite), except for APR6, for which three spectra were detected. See **Table S1** for the complete list of proteins identified.

The apical complex contains a cytoskeletal scaffold as well as associated membrane-bound secretory organelles [5]. In *Toxoplasma,* the cytoskeletal apical complex is well characterized structurally for mature parasites. The major elements are the conoid, the preconoidal rings, and intra-conoid microtubules (MTs), as well as the apical polar ring, which associates with the minus ends of 22 evenly-spaced cortical MTs [6-12]. Components of the cytoskeletal apical complex are involved in signaling and the mechanics of parasite invasion into the host cell [13-22]. The conoid is made of ribbon-like fibers, polymerized from the same tubulin subunits as the cortical MTs despite their radical structural differences. It is a mobile structure. In response to changes in the intra-parasite calcium concentration, the conoid, together with the preconoidal rings and intra-conoid MTs, protrudes or retracts through the apical polar ring [6-8]. A number of apical polar ring proteins have been identified and characterized previously [16, 17, 23-29]. These proteins individually have varying degrees of impact on different aspects of the parasite lytic cycle. Some proteins, such as KinesinA and APR1, also have a notable structural role [17]. Without KinesinA, the electron dense, well-defined annulus conventionally noted as the apical polar ring becomes undetectable by negative staining electron microscopy (EM). The removal of APR1 results in frequent conoid detachment. Interestingly, the cortical MT array appears to be normal in parasites lacking only KinesinA or only APR1. However, when both KinesinA and APR1 are knocked out, cortical MTs often detach in groups from the apex in adult parasites as well as in large daughter parasites [17, 30].

The structural integration of the apical polar ring and cortical MTs suggests a role of the apical polar ring as the organizing center for the cortical MTs. Using APR2—an early component of the apical polar ring—as the marker, we examined how the apical polar ring is constructed with respect to the cortical MT array [26]. We established that the assembly of the apical polar ring initiates prior to that of the MT array, starting from an arc with its open side facing the centrioles. As the arc grows into a ring, the MT array begins to assemble on the closed side. This observation supports the hypothesis that the apical polar ring initiates and templates new MT arrays by directional, stepwise assembly coupled with depositing or recruiting factors for polymerization of new MTs.

In this work, we use APR2 as the bait in immunoprecipitation and identified several new components of the apical polar ring. Among them is APR9, a protein found not only in apicomplexans, but also in *Chromera velia*, a free-living relative of apicomplexans. The knockout of APR9 alone results in a mild phenotype in the parasite lytic cycle. In contrast, when APR9 is removed together with KinesinA, the lytic cycle is greatly inhibited. The *ΔkinesinAΔapr9* parasite is nearly completely paralyzed, which severely impairs egress and invasion. Even though both APR9 and Kinesin A are early components of the apical polar ring, only a small fraction of the *ΔkinesinAΔapr9* parasites display perturbed MT organization. However, the *ΔkinesinAΔapr9* parasite does display abnormalities in multiple subcellular processes, including the formation of an apical actin concentration, impaired conoid extension, and significantly reduced secretion of a major micronemal adhesin (MIC2) upon stimulation with calcium ionophore. These findings reveal that, in addition to its role as an MT organizing center, the apical polar ring is involved in distinct subcellular processes and plays a critical role in parasite motility. They also highlight the importance of assessing the function of the components of the apical polar ring not only individually but also in combination with others.

## RESULTS

### Enrichment and identification of new apical polar ring proteins using APR2-mEmeraldFP as the bait

We previously showed that APR2 (TgGT1_227000) is an early component of the apical polar ring [26]. To enrich and identify new apical polar ring components, we carried out immunoprecipitation (IP) using an anti-GFP nanobody and mEmeraldFP-APR2 knock-in parasites. The parental parasite having untagged APR2 was used as the negative control. Parasites were homogenized after Triton X-100 (TX-100) extraction, and a high-affinity llama anti-GFP antibody conjugated to agarose beads ("GFP-trap") was used for the pull-down. The IPs were analyzed by <u>Mu</u>lti<u>d</u>imensional <u>P</u>rotein Identification <u>T</u>echnology (MudPIT) to identify enriched proteins [13, 31-33] (**Fig 1B**). In addition to multiple known proteins of the apical polar ring, three new candidates were also identified: TgGT1_223790 (20 <u>P</u>eptides), 289990 (5 P), and 295420 (10 P) (**Fig 1C, Table S1**). To examine the localization of these proteins, we generated knock-in lines, in which the genomic locus of the target gene is replaced with a DNA fragment including the CDS for mEmeraldFP-geneX with a 3’UTR and an expression cassette for the HXGPRT selectable marker (**Fig 2A-B**). Expansion microscopy revealed that these three proteins are localized to an apical annulus immediately above the conoid in intracellular parasites, which is the expected location for the apical polar ring (**Fig 2C-E**, insets). They are thus named APR9 (223790), APR10 (289990) and APR11 (295420).

**Fig. 2.**
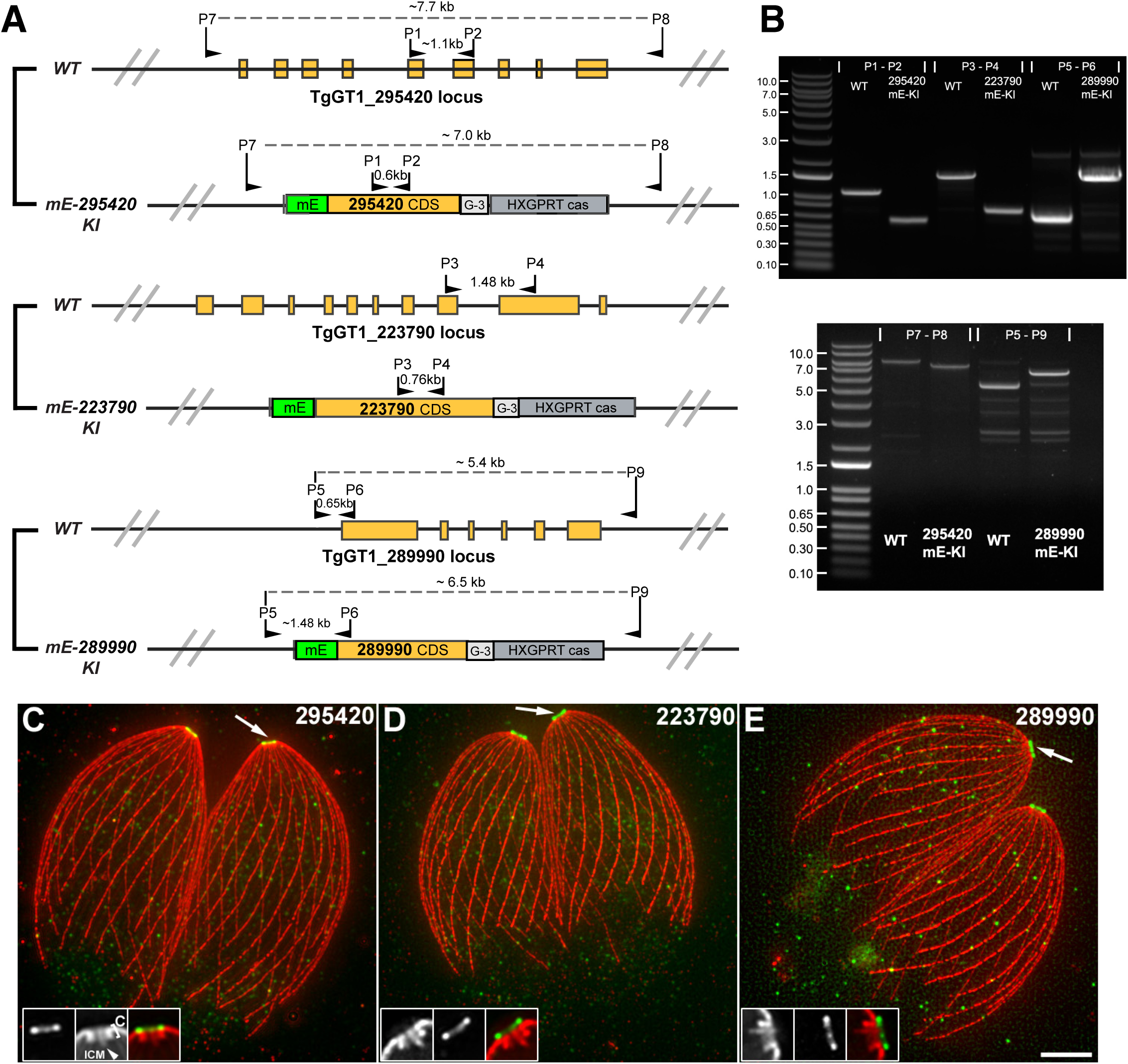
Confirmation of the localization of TgGT1_295420, 223790, 289990 to the apical polar ring. **A-B**. Generation of mEmeraldFP tagged 295420, 223790, 289990 knock-in parasites. A: Schematic illustrating the strategy used to generate the mEmeraldFP-tagged knock-in parasites. Positions for the primers in the diagnostic genomic PCRs shown in B and expected DNA fragment sizes are indicated. B: Diagnostic genomic PCRs of the RH*Δku80* parental (WT) and the knock-in lines confirming the homologous integration of the mEmerald fusion in the knock-in lines. **C-E.** Projections of expansion microscopy (ExM) images of mEmeraldFP-tagged 295420, 223790, 289990 knock-in parasites labeled with anti-GFP and anti-tubulin antibodies. Insets (2X) correspond to the regions indicated by the arrows, which show that TgGT1_295420, 223790, 289990 are localized to an apical annulus positioned anterior to the retracted conoid (C, bracket), consistent with the localization of the apical polar ring. Arrowhead indicates intra-conoid MTs (ICM). Scale bar indicates an estimated length of ∼ 1 μm before expansion and is based on an estimated expansion ratio of ∼5.4 [26, 30]. Image contrast was adjusted to optimize display.

Of the three new APR proteins, we decided to first focus on APR9. Aside from weak homology with the ENBA-3A domain at its C-terminus, APR9 does not have any readily recognizable protein domains. However, APR9 is highly conserved within Apicomplexa, and is found in all sequenced apicomplexan lineages. Furthermore, a well-conserved homolog (Cvel_27126) is also found in *Chromera*, a free-living relative of the apicomplexans (**Fig 3A**). In *T. gondii*, APR9 shares strong sequence similarities with Tg219500, which has previously been localized to the apical polar ring and named as APR4 in [27]. Multiple APR9/4 homologs are found in the genomes of members of the Sarcocystidae family (*e.g*., *T. gondii, Neospora caninum* and *Sarcocystis neurona*), likely a result of a gene duplication event in their common ancestor. Alpha-fold 3 [34] predicts that both APR4 and APR9 have a core of multiple alpha-helices, although TgAPR9 and its orthologs in *Plasmodium berghei* and *Chromera velia* are predicted to contain extended unstructured regions (**Fig S1**).

**Fig. 3.**
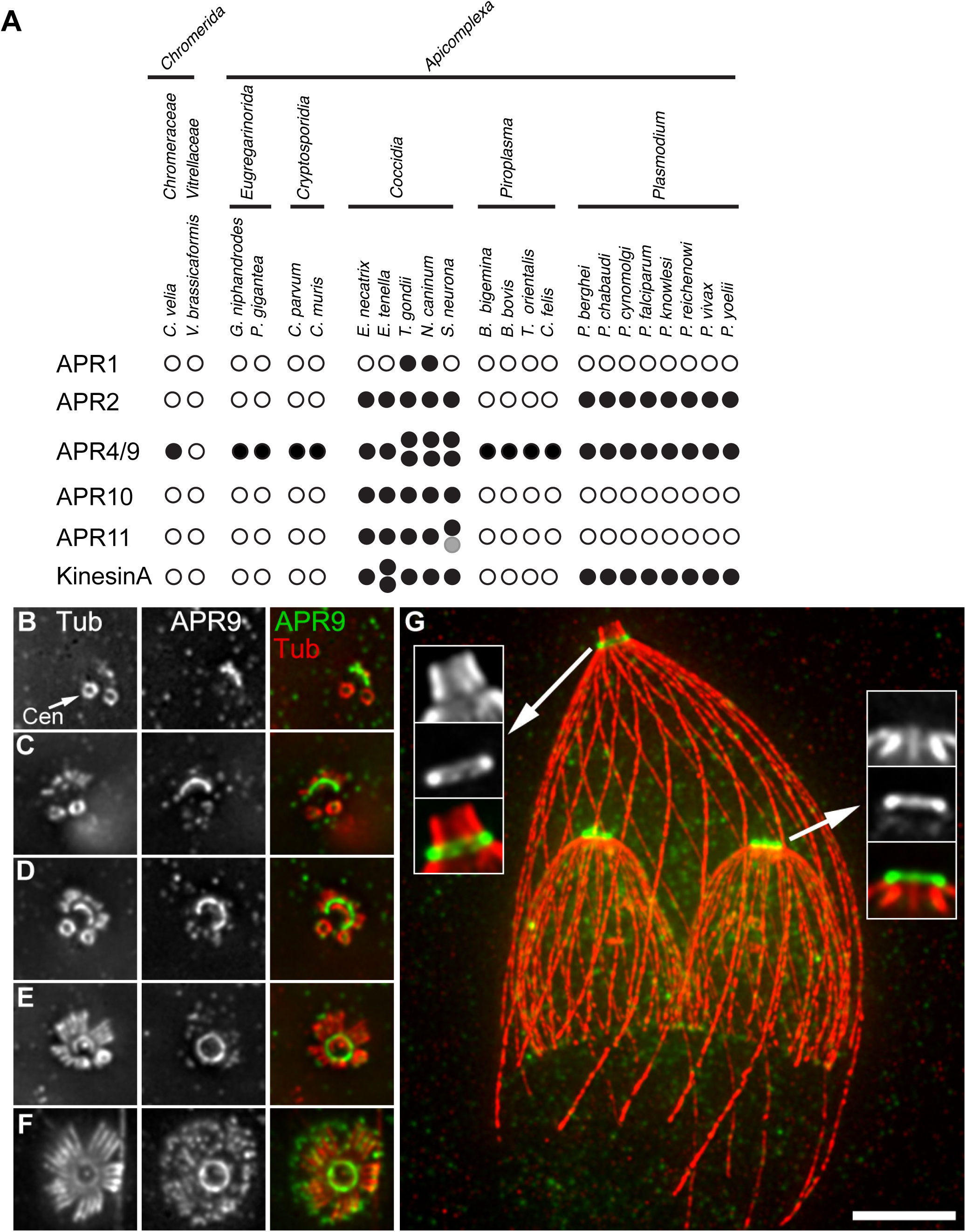
APR9 is broadly conserved among the apicomplexans and their free-living relative *Chromera velia,* and is recruited to the apical polar ring early during daughter assembly. **A.** Presence (filled circle) or absence (open circle) of predicted homologs for TgAPR1, APR2, APR9, APR10 and APR11 and KinesinA in representative members of the phylum Apicomplexa with available genome sequences, as well as their photosynthetic relatives, *Chromera velia* and *Vitrella brassicaformis*. Protein sequences of *T. gondii* homologs were used as queries for the BLAST searches against the predicted protein databases in VEupathDB (http://veupathdb.org). An E-value cut-off of 1e-5 was used to identify APR1, APR2, APR9, APR10, and APR11 orthologs. Grey circle indicates a hit with 1e-5> E-value > 1e-10. Due to the conserved nature of the kinesin-fold, only hits with an E-value less than 1e-16 were considered tor KinesinA orthologs, and were manually curated. **B-G.** Projections of ExM images of *mE-APR9* knock-in parasites through daughter development. Insets (2X) in G show that the APR9 labeling is apical to the retracted daughter conoid (single section),but posterior to the protruded conoid in the mother parasite (projection of a substack). Cen: centrioles. Scale bar ≈ 1 µm prior to expansion based on an estimated expansion ratio of ∼5.4 [26, 30]. Image contrast was adjusted to optimize display.

### APR9 is an early component of the apical polar ring and its localization is independent of APR2, APR4 and Kinesin A

To determine the recruitment schedule of APR9 in developing daughters, we used expansion microscopy and labeled the FP-knock-in parasite with anti-GFP and anti-tubulin antibodies. The timing of recruitment of APR9 was determined by using the number and length of cortical MTs to estimate daughter developmental stages. Similar to APR2, APR9 is assembled into the precursor of the apical polar ring prior to the appearance of the daughter MTs. As reported before [26], the apical polar ring extends towards the centriole region from an arc to a closed ring. The APR9 signal in the apical polar ring persists throughout the daughter development and in the mature parasite (**Fig 3B-G**). In addition to the prominent localization to the apical polar ring, there is also notable mE-APR9 signal in the daughter cortex (**Fig 3F**). To determine how the localization of APR9 is affected by other known apical polar ring components, we generated *mEmerald* (*mE*)-*APR9* knock-in lines in the *Δapr2, Δapr4,* and *ΔkinesinA* parasites (**Fig 4A-E, Fig S2**). APR9 is targeted to an apical annulus in these lines, indicating that the localization of APR9 is independent of APR2, APR4, and KinesinA (**Fig 4C-E**). Interestingly, while the positioning of the APR9 ring appears to be normal in the *Δapr2* and *Δapr4* parasites, it occasionally appears to be tilted in the *ΔkinesinA* parasite (**Fig 4E**, insets).

**Fig. 4.**
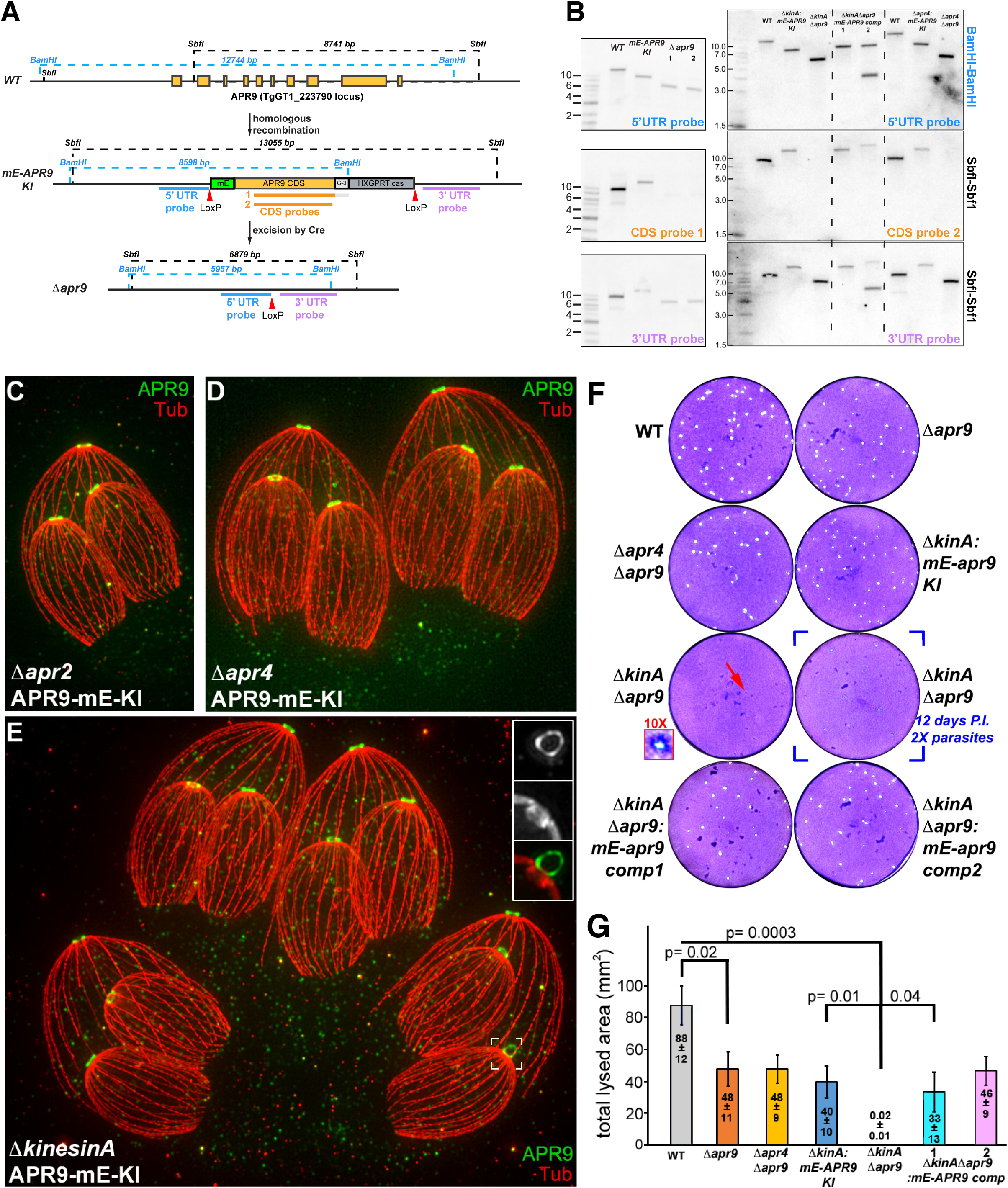
APR9 moderately affects the parasite lytic cycle when removed alone or together with APR4, but displays a strong synergistic effect with KinesinA. **A-B.** Southern blot analysis of the *apr9* locus in the WT, *mEmeraldFP- APR9* (mE-APR9 KI)*, Δapr9, ΔkinesinA*:mE-APR9 knock-in (*ΔkinA*:mE-APR9 KI)*, ΔkinesinAΔapr9* (*ΔkinAΔapr9*)*, ΔkinesinAΔapr9*:mE-APR9 complement clone 1 and clone 2*, Δapr4*:mE-APR9 knock-in, and *Δapr4Δapr9* parasites. A: Schematic for the predicted APR9 locus in WT, *mE-*APR9 knock-in, and *Δapr9* lines. Restriction sites, hybridization targets of the Southern blot probes for the *apr9* coding region (CDS probe 1 and 2, orange bars), regions upstream (5’ UTR probe, blue bar) and downstream (3’ UTR probe, purple bar) of the CDS, and the expected DNA fragment sizes are indicated. B: Southern blot analysis confirming the homologous integration of the mE-APR9 fusion in the knock-in lines, and the deletion of the *apr9* locus in the *Δapr9* lines. See Fig. S2 for Southern blot analysis of the *apr4* locus in the APR4 knock-in and knockout lines. **C-E.** Projections of expansion microscopy (ExM) images of *mE-APR9* knock-in in *Δapr2* (C)*, Δapr4* (D) and *ΔkinesinA* (E) parasites. Inset (2X) in E indicates a daughter APR9 ring tilted relative to the parasite cortex. **F.** Plaque assays of WT, *Δapr9*, *Δapr4Δapr9, ΔkinesinA*:mE-APR9 knock-in (*ΔkinA*:mE-APR9 KI)*, ΔkinesinAΔapr9* (*ΔkinAΔapr9*), and *ΔkinesinAΔapr9*:mE-APR9 complement clone 1 and clone 2 (*ΔkinAΔapr9*:mE-APR9 Comp 1 and 2) parasites. All cultures except for the bracketed condition were infected with 100 parasites and incubated for 7 days before crystal violet staining. The bracketed culture was infected with twice as many Δ*kinA*Δ*apr9* parasites and incubated for 5 more days, yet still produced very few and very small plaques. Inset (10X) highlights the small plaque indicated by the red arrow. **G.** Bar graphs that quantify plaquing efficiency for the lines shown in F. Three to seven independent plaque assays were performed for each line. For each replicate, the total lysed area of three wells, each infected with 100 parasites for 7 days, was measured. Error bars represent standard error of the mean (SEM). Value indicated in each bar is average ± SEM for the corresponding line. P-values were calculated in KaleidaGraph using two-tailed unpaired Student’s *t*-tests with unequal variances.

### Impact of APR9 deletion on the parasite lytic cycle

To determine the impact of APR9 on the parasite lytic cycle, we generated a knockout line, *Δapr9,* by transient expression of Cre-recombinase in the *mE-APR9* knock-in parasite to excise the LoxP flanked region (**Fig 4A-B**). Putative knockout clones were first identified by the loss of mE-APR9 fluorescence. The excision of the mE-APR9 +selectable marker in the LoxP-flanked region was then confirmed by Southern blotting. Parasite growth was assessed by the ability of parasites to generate "plaques" over 7 days of incubation with host cell monolayers. The plaques are lesions in the monolayer generated by continuing destruction of the host cells by the parasite via cycles of invasion, replication, and egress. The *Δapr9* parasite shows only moderate growth defects in plaque assays (**Fig 4F-G**). This is consistent with the prediction from a previous genome-wide CRISPR-Cas9 screen, which assigned a phenotype score of 0 for APR9 (ToxoDB.org, [35], **Fig 1**).

Given the broad conservation of APR9 which extends to a free-living relative of apicomplexans, the weak phenotype of the *Δapr9* parasite is somewhat surprising. One possible explanation is the function and/or structural redundancy among the apical polar ring components. Because APR4 is a close homolog of APR9, we generated a double-knockout (*Δapr4Δapr9*) by transiently expressing the Cre-recombinase in the *Δapr4*:*mE-APR9* knock-in parasite to excise the mE-*APR9* knock-in locus (**Fig 4A-B**). We similarly generated a *ΔkinesinAΔapr9* double knockout line because KinesinA shows a synergistic effect with two other apical polar components, APR1 and APR2 [17, 26]. We found that the *Δapr4Δapr9* parasite shows only a moderate defect in plaque assays, similar to the *Δapr9* parasite (**Fig 4F-G**). In contrast, the phenotype of the *ΔkinesinAΔapr9* parasite is severe. The *ΔkinesinAΔapr9* clones were isolated with great difficulty. After 7 days of incubation for the plaque assay, the parental lines formed many sizable plaques, but the few plaques in the *ΔkinesinAΔapr9* culture were barely visible (**Fig 4F**). The plaquing efficiency of the *ΔkinesinAΔapr9* parasite is ∼ 0.03% of the wild-type parasite, ∼ 0.05% of *Δapr9*, and ∼ 0.06% of *ΔkinesinA*:*mE-APR9* knock-in parasite, its immediate parental line (**Fig 4G**). We generated two complemented lines (*ΔkinesinAΔapr9*:mE-APR9 complement clone 1 and clone 2) in which the mE-APR9 expression cassette is integrated in different genomic loci in the *ΔkinesinAΔapr9* parasite (**Fig 4B**). Plaquing efficiencies in both complemented lines are restored to close to the level of the *ΔkinesinA*:*mE-APR9* knock-in parental line (**Fig 4F-G**).

### The loss of KinesinA and APR9 slows parasite replication and severely compromises parasite motility

To further determine the specific defects in the *ΔkinesinAΔapr9* parasite, we examined its replication, invasion, and egress behaviors. We found that the *ΔkinesinAΔapr9* parasites replicate with a doubling rate lower than that of the WT parasites (**Fig 5A**). The replication rate of the WT parasite between 12 hr and 36 hr is 2.95 doublings/24hr. The doubling of the *ΔkinesinAΔapr9* parasite is 2.2 doublings/24hr. Although this difference is statistically significant (p = 0.0025), it translates to only ∼32 fold fewer parasites over 7 days. This cannot fully explain the drastic difference in plaquing efficiency, which differs by more than 3,700 fold between the WT and the *ΔkinesinAΔapr9* parasites. Other aspects of the lytic cycle must also be affected by the loss of KinesinA and APR9. Indeed, host cell invasion of the *ΔkinesinAΔapr9* parasite is severely impaired when assessed using a dual-color invasion assay that distinguishes between intracellular and extracellular parasites based on antibody accessibility to a surface antigen, P30 [36, 37] (**Table 1**). The *ΔkinesinAΔapr9* parasite invades at ∼ 11% (p < 0.0001) of the level of the WT parasites. Both invasion and egress rely on active gliding motility of the parasite. To determine if the motility of *ΔkinesinAΔapr9* is affected, we carried out live egress assays induced by the calcium ionophore, A23187. The calcium ionophore treatment elevates the calcium concentration in the parasite cytoplasm. In the wild-type parasite, upon A23187 treatment, the combined effects of active parasite movement and host-cell-membrane permeabilization by pore-forming proteins secreted from the parasite enable a rapid dispersal of the parasites from the host cell (**Fig 5B-C**). The *ΔkinesinAΔapr9* parasites, however, were nearly completely paralyzed. Most parasites remained immotile. Only sporadic movement was observed in a small minority of the parasites (**Fig 5B-C**, Video S1). Notably, this phenotype is specific to motility but not due to a universal block of calcium sensing because the *ΔkinesinAΔapr9* parasites still respond to the A23187 treatment by secreting factors to permeabilize the host cell membrane (**Fig 5B**, Video S1). This is indicated by the abrupt change in morphology and contrast of the host cell in the DIC image as well as the rapid labeling of the host cell nucleus by the cell-impermeant dye DAPI upon A23187 treatment.

**Fig. 5.**
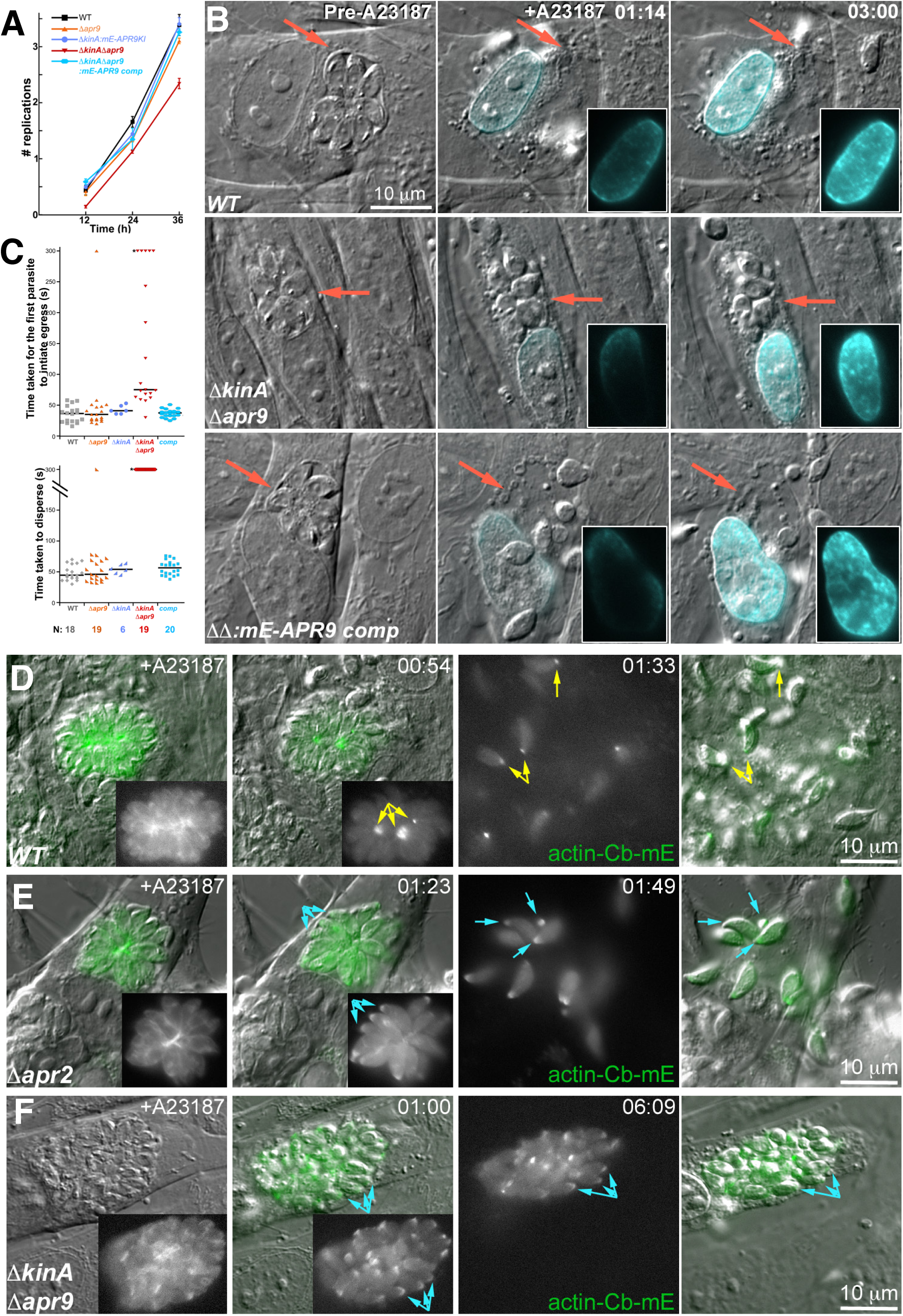
Removal of APR9 and Kinesin A together slows parasite replication, drastically reduces motility, and disrupts actin dynamics. **A.** Replication assay of RH*Δku80* parental (WT), *Δapr9*, *ΔkinesinA*:*mE-APR9* knock-in (*ΔkinA*:*mE-APR9* KI)*, ΔkinesinAΔapr9* (*ΔkinAΔapr9*), and *ΔkinesinAΔapr9*:*mE-APR9* complement parasites. X-axis: time after the start of parasite-host cell incubation. Y-axis: number of parasite doublings. Error bars: SEM. **B.** Time-lapse images of A23187-induced egress of RH*Δku80* parental (WT), *ΔkinesinAΔapr9* (*ΔkinAΔapr9*), and *ΔkinesinAΔapr9*:*mE-APR9* complement (*ΔΔ*:*mE-APR9* comp) parasites. The cell-impermeant nucleic acid dye DAPI was added to the medium prior to the A23187 treatment. Labeling of the host cell nuclear DNA indicates host-cell permeabilization. Note that although *ΔkinAΔapr9* parasites display drastically reduced motility, they still secrete factors to lyse the host cell upon A23187 treatment, as indicated by the change in the host-cell morphology in the DIC images and by DAPI entering and binding to DNA in the host cell nucleus (Video S1). Insets (1X) show DAPI images of the host cell nucleus, with contrast adjusted to clearly display labeling at the rim of the nucleus. **C.** Dot plots of time taken for the first parasite to initiate egress and time taken to disperse during A23187-induced egress for RH*Δku80* parental (WT), *Δapr9*, *ΔkinesinA, ΔkinesinAΔapr9*, and *ΔkinesinAΔapr9*:*mE-APR9* complement (comp) parasites. The median for each dataset is indicated by a black bar. Time-taken-to-disperse for each vacuole was defined as the time point at which more than 50% parasites have actively egressed. *ΔkinesinAΔapr9* parasites did not reach this threshold in any vacuole during the 300-second observation period (*). **D-F.** Live imaging of RH*Δku80* parental (WT, D), *Δapr2* (E), and *ΔkinesinAΔapr9* (F) parasites expressing actin-Cb-mE and treated with 5 µM A23187. Yellow arrows: basal actin-Cb-ME accumulation. Cyan arrows: apical actin-Cb-mE accumulation.

**Table 1.** Quantification of invasion for five *T. gondii* lines. For each line, the number of intracellular parasites per field was counted in 10 fields in each of five independent biological replicates. SEM.: Standard error of the mean. P-values for "%WT" are indicated on the right and were calculated in KaleidaGraph using two-tailed unpaired Student’s *t*-tests with unequal variance. In total (intracellular + extracellular), 11,795 parasites were counted for the RH*Δku80* parental line, 12,692 for *ΔkinesinA*:mE-APR9 knock-in, 11,684 for *Δapr9*, 10,575 for *ΔkinesinAΔapr9*, and 16,055 for *ΔkinesinAΔapr9:APR9* complement parasites.

| Strain | Number invaded ± SEM | % WT ± SEM | RH*Δku80* (WT) | *ΔkinesinA:mE-APR9* knock-in | *Δapr9* | *ΔkinesinAΔapr9* | *ΔkinesinAΔapr9:APR9* complement |
| --- | --- | --- | --- | --- | --- | --- | --- |
| RH*Δku80* (WT) | 64±27 | 100 |  | p= 0.37 | p= 0.98 | **p<0.0001** | p= 0.22 |
| *ΔkinesinA:mE-APR9* knock-in | 48±21 | 85±17 |  |  | p= 0.67 | **p=0.007** | p=0.59 |
| *Δapr9* | 42±10 | 99±32 |  |  |  | **p= 0.038** | p= 0.45 |
| *ΔkinesinAΔapr9* | 5±2 | 11±3 |  |  |  |  | **p= 0.04** |
| *ΔkinesinAΔapr9:APR9* complement | 32±12 | 71±23 |  |  |  |  |  |

### The loss of KinesinA and APR9 results in an accumulation of actin at the apical portion of the parasite upon stimulation with a calcium ionophore

Parasitic gliding motility is dependent on actin polymerization and associated myosins. The current model predicts an apical-basal actin flux powered by cortex-associated myosins [38]. In recent years, an mEmerald-tagged actin nanobody ("actin-chromobody") that preferentially associates with F-actin has been used to assess actin kinetics [18, 39-41]. The most notable observation is the buildup of a basal accumulation of mEmerald fluorescence, presumed to be actin-nanobody complexes when the parasite motility is stimulated. To determine how the loss of KinesinA and APR9 affects actin kinetics, we transiently expressed the mEmerald-actin-chromobody (mE-actin-Cb) in WT and *ΔkinesinAΔapr9* parasites and treated intracellular parasites with A23187 to induce egress (**Fig 5D-F**). In contrast to the WT parasite, where mE-actin-Cb builds up at the basal end of the parasite upon A23187 treatment, mE-actin-Cb accumulates in an apical cap in the *ΔkinesinAΔapr9* parasite. Previously it was shown that the knockdown of APR2 results in apical mE-actin-Cb concentration when treated with BIPPO, an inhibitor of apicomplexan phosphodiesterases implicated in the Ca^2+^ signaling pathways [27, 42]. We also found the formation of a prominent apical mE-actin-Cb concentration when our APR2 knockout line (*Δapr2*) was treated with A23187. However, as we reported before [26], the *Δapr2* parasites actively move out of the host cell during A23187-induced egress (**Fig 5E**). This indicates that the buildup of an apical actin concentration does not have a major impact on parasite motility, and cannot explain the motility defect in the *ΔkinesinAΔapr9* parasites.

### The loss of KinesinA and APR9 only has a minor impact on the organization of cortical MTs, but severely compromises conoid protrusion

Although APR9 and KinesinA are both early components of the apical polar ring, the loss of KinesinA and APR9 does not have a major impact on the organization of the cortical MTs (**Fig 6A**). Only ∼15% of mature intracellular *ΔkinesinAΔapr9* parasites have detached MTs or abnormal MT organization (total 177 parasites counted). To determine how APR9 and KinesinA impact the overall structure of the apical complex, we carried out negative staining EM analysis of TX-100 extracted parasites. Similar to the *ΔkinesinA* parasite [17], *ΔkinesinAΔapr9* does not have a strongly stained annulus at the apical ends of the cortical MTs. There is no overt structural abnormality in the conoid of *ΔkinesinAΔapr9* parasites (**Fig 6B**). The conoid is a motile organelle and extends upon calcium stimulation through the apical polar ring. To determine whether conoid protrusion is affected in the *ΔkinesinAΔapr9* parasite and its parental lines, we treated extracellular parasites with A23187 for 1 min, fixed with formaldehyde and processed for expansion microscopy labeled with anti-tubulin antibody (**Fig 6C**). While over ∼96% of the wild-type parasites were found with prominent, extended conoid (total 198 parasite counted), only ∼ 31% of the *ΔkinesinAΔapr9* parasites have a fully protruded conoid (total 249 parasites counted) (**Fig 6C-D**). The conoid and the associated preconoidal rings contain multiple motility-related signal and structural proteins [14, 15, 18, 19]. It is thus conceivable that blocking the movement of this structure loaded with motility-relevant factors could globally impact parasite motility. However, we found that only ∼10% of the *Δapr2* parasites fully extend the conoid (total 165 parasites counted) (**Fig 6D**), even though this knockout moved actively and directionally during A23187-induced egress (**Fig 5E** and [26]). The lack of conoid extension under this condition therefore cannot, by itself, explain the motility phenotype of the *ΔkinesinAΔapr9* parasites.

**Fig. 6.**
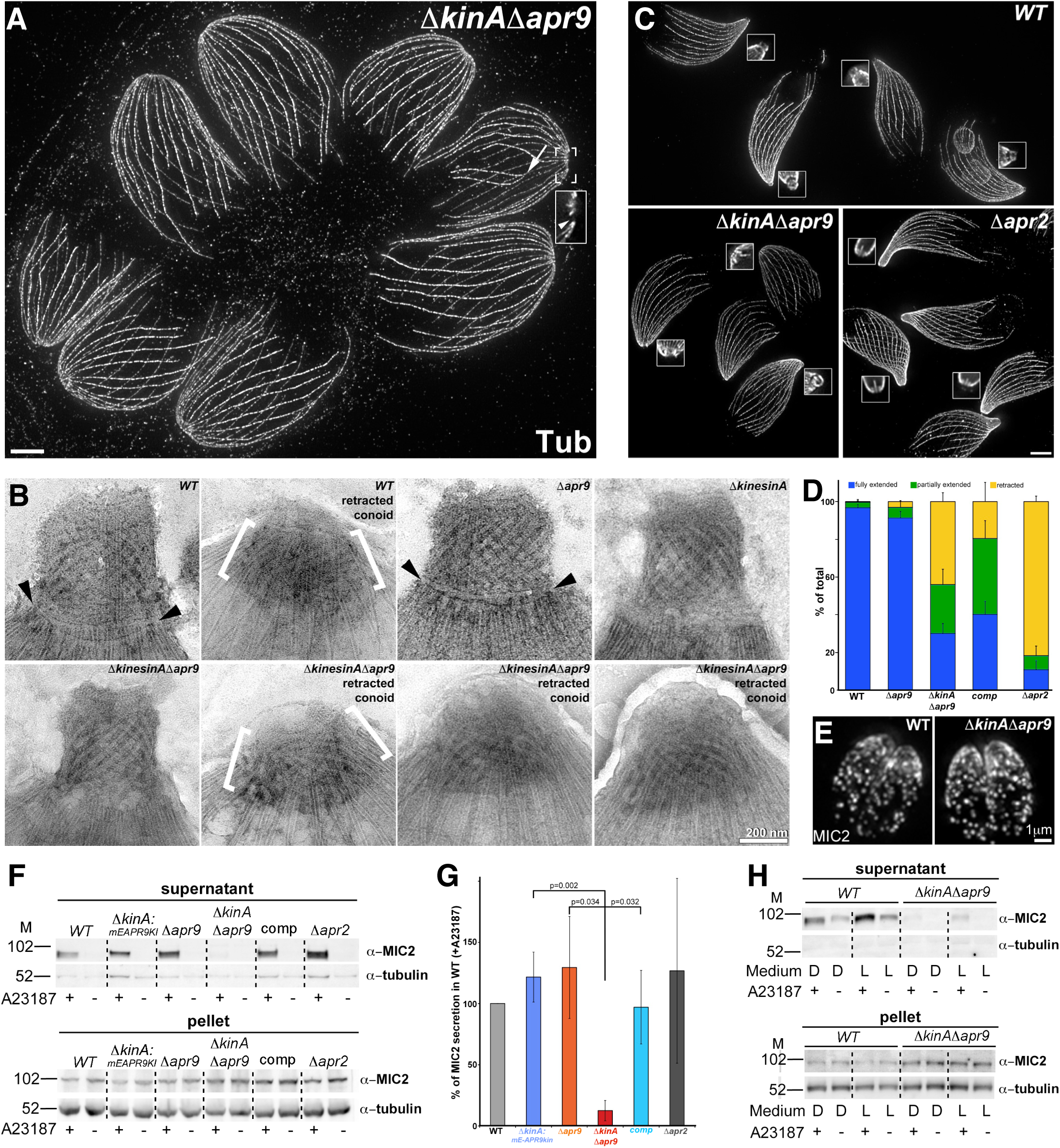
The impact of the loss of KinesinA and APR9 on the organization of cortical MTs, conoid orotrusion, and micronemal secretion. **A.** Projections of ExM images of eight intracellular *ΔkinesinAΔapr9* parasites labeled with an anti-tubulin antibody. Cortical MT organization appears normal in most parasites. Arrow and arrowhead in the 1.5X inset indicate cortical MTs detached from the apex in one parasite. **B.** Negative-staining EM images of TX-100 extracted RH*Δku80* parental (WT)*, Δapr9, ΔkinesinA,* and *ΔkinesinAΔapr9* parasites. Arrowheads in the WT and *Δapr9* images indicate the well-defined annulus, a readily recognizable feature of the apical polar ring. This annulus is undetectable in *ΔkinesinA* and *ΔkinesinAΔapr9* parasites, although the cortical MTs remain converged at the parasite apex. For WT and *ΔkinesinAΔapr9* parasites, examples with both protruded and retracted conoid are shown. Two retracted conoids are indicated by brackets. **C.** Projections of ExM images of extracellular RH*Δku80* parental (WT), *ΔkinesinAΔapr9,* and *Δapr2* parasites after treated with 5 µM A23187 for 1 min and labeled with an anti-tubulin antibody. **D.** Bar graphs that quantify percentage of extended (blue), partially extended (green), and retracted (yellow) conoids in the extracellular RH*Δku80* parental (WT), *Δapr9, ΔkinesinAΔapr9, ΔkinesinAΔapr9:APR9* complement (comp) and *Δapr2* parasites after treatment with 5 µM A23187 for 1 min. **E.** Anti-MIC2 immunofluorescence labeling of intracellular RH*Δku80* parental (WT) and *ΔkinesinAΔapr9* parasites. **F.** Western blot analysis of the secreted and pellet fractions of RH*Δku80* parental (WT), *ΔkinesinA*:*mE-APR9* knock-in*, Δapr9*, *ΔkinesinAΔapr9* (*ΔkinAΔapr9*)*, ΔkinesinAΔapr9*:mE-APR9 complement (comp), and *Δapr2* parasites with (+) or without (-) A23187 treatment in DMEM growth medium. MIC2 was detected using the mouse anti-MIC2 antibody 6D10 [56]. Tubulin in the pellet fraction, detected by an anti-tubulin antibody, served as the loading control. M: molecular weight markers (kDa). Image contrast was inverted and enhanced to facilitate visualization. **G.** Quantification of the experiments shown in F. Values represent background-subtracted intensities of the secreted MIC2 bands in the A23187-induced samples relative to WT, normalized to the corresponding tubulin loading control in the pellet fraction. Negative background-subtracted values were set to zero. Three independent biological replicates were carried out for *Δapr2* parasite, and five for the other lines. **H.** Western blot analysis of the secreted and pellet fractions of RH*Δku80* parental (WT) and *ΔkinesinAΔapr9* parasites with (+) or without (-) A23187 treatment in DMEM growth medium (D) and L15 imaging medium (L). See **Table S2** for quantification.

### The loss of KinesinA and APR9 results in significantly reduced secretion of the major micronemal adesin, MIC2

In order for the parasite to glide, the internal force generated by myosins traveling on the polymerizing F-actin needs to be relayed by associated proteins through transmembrane adhesins. The resulting cortical force powers the parasite gliding on a surface. The major adhesin secreted by the parasite is the micronemal protein MIC2, which forms a complex with M2AP to mediate parasite gliding during invasion and egress [43-46]. The localization of MIC2 appears to be normal in intracellular *ΔkinesinAΔapr9* parasites (**Fig 6E**). We then examined MIC2 secretion with and without A23187 treatment by Western blot (**Fig 6F-H**). We first tested MIC2 secretion in DMEM parasite growth media (**Fig 6F-G**). Under this condition, the parental lines (WT, *Δapr9*, *ΔkinesinA*:mE-APR9 knock-in) and the *ΔkinesinAΔapr9*:mE-APR9 complement displayed robust MIC2 secretion upon A23187 treatment. In contrast, MIC2 secretion from the *ΔkinesinAΔapr9* parasites was much reduced. Because the egress assays revealing the profound motility defect of the *ΔkinesinAΔapr9* parasites was carried out in an L15-based imaging medium, we also examined MIC2 secretion of the *ΔkinesinAΔapr9* and WT parasites in this medium. Although the secretion of *ΔkinesinAΔapr9* parasites remained significantly lower than that of the WT parasite, both parasite lines secreted more robustly in L15 imaging medium (**Fig 6H**, **Table S2**). Previous reports showed that the complete removal of MIC2 results in a major defect in active dispersion during A23187-induced egress [46]. However, even ∼ 5% of WT MIC2 expression can support active motility during induced egress [45]. Therefore, factors other than or in addition to reduced bulk MIC-secretion underlie the motility phenotype of the *ΔkinesinAΔapr9* parasites during A23187-induced egress.

## DISCUSSION

The apical complex is an ancestral feature of the apicomplexans. It is shared by not only the thousands of apicomplexan parasites, but also their free-living relatives, such as the chromerids and colpodellids. Structural details of the cytoskeletal apical complex, such as the presence or absence of a complete conoid, varies across different apicomplexan lineages [5, 47]. However, an apical polar ring coupled with cortical MTs is found in all apicomplexans for which ultrastructural data is available. The conserved nature of the apical polar ring and its relevance to multiple aspects of parasite biology underlie the need for a deeper understanding of its composition and function.

Work in *Toxoplasma gondii* so far has identified more than a dozen components of the apical polar ring [16, 17, 23-29]. The probable extensive and complex interaction among these components is for the most part still unknown. For instance, we used APR2 as the bait in our immunoprecipitation protocol that led to the identification of APR9, yet the localization of APR9 is not perturbed in parasite lacking APR2. No doubt many other examples of these second order association will be found as more apical polar ring components are identified.

Interestingly, the known components of the apical polar ring are often poorly conserved outside Coccidia (**Fig 3A**). APR9, on the other hand, is not only broadly conserved among the apicomplexans, but also found in the free-living *Chromera velia.* Additionally, it has a close homolog, APR4, in *Toxoplasma.* The *P. berghei* ortholog of APR9 is predicted to be highly fitness-conferring (denoted “essential” in [48] and *PlasmoDB*). However, in *Toxoplasma,* the *Δapr9, Δapr4,* and *Δapr4Δapr9* parasites all have only minor phenotypic defects (this work and [27]). One might wonder under what conditions these conserved proteins perform functions sufficiently beneficial to the parasites to necessitate their retention across a wide range of apicomplexans and related lineages. One plausible explanation is a synergistic action between the apical polar ring components. In *Toxoplasma,* this effect is particularly pronounced when an additional apical polar ring component is removed from the *ΔkinesinA* parasite. So far we have generated *ΔkinesinAΔapr1, ΔkinesinAΔapr2,* and *ΔkinesinAΔapr9* parasites. The defects of all of these double-knockouts are far more severe than if the impact of the target genes were simply additive ([17, 26] and this work).

*ΔkinesinAΔapr9* parasite displays a pronounced motility defect. Most parasites appear to be paralyzed during A23187-induced egress. At the same time, the *ΔkinesinAΔapr9* parasite develops an apical actin concentration, as visualized by mE-actin-Cb. This, however, cannot explain the motility defect in the *ΔkinesinAΔapr9* parasite, since the *Δapr2* parasites, which also exhibit the accumulation of an apical actin cap, are motile and egress normally upon A23187 treatment. The *Δapr2* and the *ΔkinesinAΔapr9* parasites also both display a defect in conoid extension. This suggests that the apical polar ring is important for the mobility of the conoid. Together with the previously reported role of the apical polar ring in conoid attachment and positioning [17, 27], multiple aspects of the structural and function dependence of these two macromolecular complexes have now emerged. Conoid protrusion, however, is not tightly coupled with parasite motility, as the *Δapr2* parasite is highly impaired in conoid extension but actively moves out of the host cell during A23187-induced egress.

To move by gliding, the parasite needs to adhere to the surface that it interacts with. MIC2 has been proposed to be the major adhesin that connects the internal actomyosin machinery with the surface that the parasite glides on. Reduced MIC2 secretion results in a significant defect in parasite invasion [45]. Previous studies revealed that several components of the apical polar ring contribute to the secretion of MIC2. The knockdown of RNG2 significantly reduced the constitutive secretion of MIC2 [16]. The loss of both KinesinA and APR1 results in a major decrease in ethanol-induced MIC2 secretion. This work shows a large reduction of A23187-induced MIC2 secretion from the *ΔkinesinAΔapr9* parasite, which likely contributes to the invasion defect of this double-knockout and reaffirms the role of the apical polar ring in MIC2 secretion. Notably, unlike in the *ΔkinesinAΔapr1* parasite [17], the organization of cortical MTs is largely unaffected in the *ΔkinesinAΔapr9* parasite. This indicates that the role of the apical polar ring in controlling MIC2 secretion is independent of its role as an MT organizing center. It is also worth noting that the bulk secretion of MIC2 from *ΔkinesinAΔapr9* parasite, although considerably reduced, is still detectable, especially in L15 imaging medium, where this double-knockout displays profound motility defects during A23187-induced egress. Given that even a small residual level of MIC2 expression can support active parasite movement during induced egress [45], we propose that the motility defect in *ΔkinesinAΔapr9* parasite is a consequence of combined effects of abnormalities in actin kinetics, conoid protrusion and micronemal secretion. Alternatively, the apical polar ring dictates another, so far unknown, aspect of motility control through KinesinA and APR9.

Interestingly, while both KinesinA and APR9 are early components of the apical polar ring, their loss perturbs microtubule organization in only a minority of the parasites. Nevertheless, the broad conservation of APR9 among the apicomplexans is of great interest, especially in the free-living *Chromera velia.* We believe that it is a useful probe for structurally defining the counterpart of the apical polar ring in *Chromera* and exploring its construction in conjunction with other apical complex-related structures, such as the pseudoconoid. The role of APR9 and other conserved motility-relevant factors (e.g. AKMT) associated with the apical complex also pose the questions regarding how the apical complex was involved in the origination of gliding motility, whether this behavior predates the development of intracellular parasitism, and if so, how it was utilized in the life of the free-living ancestor of apicomplexans, the motile form of which was likely to be flagellated like *Chromera*.

## MATERIALS AND METHODS

### T. gondii cultures

As described in [17, 19, 49, 50], *T. gondii* tachyzoites were maintained in confluent cultures of human foreskin fibroblasts (HFFs) in DMEM growth medium [Dulbecco’s Modified Eagle’s Medium (Corning, 15-013-CV) supplemented with 1% (v/v) heat-inactivated cosmic calf serum (SH30087.3; Hyclone, Logan, UT) and Glutamax (Life Technologies-Gibco, 35050061)].

### Immunoprecipitation and Multidimensional Protein Identification Technology (MudPIT) analysis

Immunoprecipitation experiments were performed as described in [51] using the mEmerald-APR2 knock-in parasites [26]. Untagged RH*ΔhxΔku80* parasites (a kind gift from Dr. Vern Carruthers at the University of Michigan [52]) was used as the negative control. Parasites were processed as described in [51], except that the lysis buffer contained 1% TX-100 instead of 0.5%. The lysate was clarified by centrifugation and then incubated with Chromotek-GFP-Trap agarose beads (AB_2631357, Proteintech) for 90 minutes at 4°C before elution for MudPIT analysis.

Protein samples were processed for MudPIT and analyzed as described in [19, 51]. Raw data and search results files have been deposited to the Proteome Xchange (accession: PXD074785) via the MassIVE repository and may be accessed at ftp://MSV000100958@massive-ftp.ucsd.edu with password Kehu-2026-02-23. Original mass spectrometry data underlying this manuscript can be accessed after publication from the Stowers Original Data Repository at http://www.stowers.org/research/publications/libpb-2610.

*Plasmid construction* (See **Table S3** for primers used in this study)

Genomic DNA (gDNA) and coding sequences (CDS) were prepared as described in [19]. *pTKO2_II-mEmerald-APR4 (mE-APR4)* knock-in plasmid: ∼1.9 kb sequences upstream (5’UTR) or downstream (3’UTR) of the APR4 (TgGT1_ 219500) genomic locus were amplified from the parasite gDNA by PCR using primer pairs S1/ AS1 and S2/AS2, respectively, and inserted at the *Not*I (5’UTR) or *Hind*III (3’UTR) site of plasmid pTKO2-II-mCherryFP [51] using the NEBuilder HiFi Assembly kit. The coding sequences for mEmerald fluorescent protein (mEmerald) and APR4 were amplified using primer pairs S3/AS3 and S4/AS4, respectively, and assembled into the *AsiS*I site to generate pTKO2_II-mE-APR4. A five-amino acid linker (SGLRS) was inserted between the APR4 and mEmerald coding sequences, and the Kozak sequence from the endogenous APR4 locus (CCAATGG) was added to the 5’ end of the mEmerald coding sequence.

*pTKO2_II- mE-APR9*, *pTKO2_II- mE-APR10,* and *pTKO2_II- mE-APR11* knock-in plasmids: These plasmids were constructed using the same strategy as described for pTKO2_II-mE-APR4 knock-in using the primers listed in **Table S3**. The endogenous Kozak sequences added to the 5’ end of the mEmerald coding sequence were ACCATGA, GCGATGG, and GCGATGA for APR9 (TgGT1_ 223790), APR10 (TgGT1_ 289990), and APR11 (TgGT1_ 295420), respectively.

### Generation of knock-in and knockout parasite lines

*mE-APR4, mE-APR9, mE-APR10 and mE-APR11* knock-in lines: All knock-in and knockout lines were generated in the RH*ΔhxΔku80* strain, referred to throughout this study "RH*Δku80",* "wild-type" or "WT". ∼1 x 10^7^ RH*ΔhxΔku80* parasites were electroporated with 40 µg of the knock-in plasmid linearized by *Not*I, using the settings described previously [17, 19, 49]. Transfected populations were then selected with 25 µg/mL mycophenolic acid and 50 µg/mL xanthine. Because the backbone of the pTKO2_II plasmid contains a cassette driving cytoplasmic expression of mCherryFP [51], clones were first selected by positive mEmeraldFP fluorescence at the apical polar ring as well as absence of cytoplasmic mCherry fluorescence, allowing exclusion of non-homologous or single crossover recombinants. Correct homologous integrations were subsequently verified by Southern blot (for the mE-APR4 and mE-APR9 knock-in lines) or by genomic PCRs (for the mE-APR10 and mE-APR11 knock-in lines).

*Δapr4* and *Δapr9* lines:*mE*-*APR4 or mE-APR9* knock-in clones confirmed by Southern blot were electroporated with 30 µg of pmin-Cre-eGFP_Gra-mCherry, and selected with 6-thioxanthine at 80 µg/mL as described previously [19, 49]. Clones that had lost the mEmerald fluorescence were first identified by microscopic screening. The deletion of the LoxP-flanked region in the selected clones was then confirmed by Southern blot.

*Δapr4:mE-APR9* knock-in and *ΔkinesinA:mE-APR9* knock-in lines: The *Δapr4* or *ΔkinesinA* parasites [17] were electroporated with the *Not*I-linearized pTKO2_II-mE-APR9 knock-in plasmid. Knock-in clones were selected and confirmed as described above for the *mE-APR9* knock-in line.

*Δapr4Δapr9* and *ΔkinesinAΔapr9* lines: *Δapr4:mE-APR9* knock-in or *ΔkinesinA:mE-APR9* knock-in parasites were electroporated with pmin-Cre-eGFP_Gra-mCherry. Clones were selected as described above for the *Δapr9* parasite.

### Southern blotting

Southern blotting was carried out as described in [19, 26, 49, 51]. To probe and detect changes at the target genomic locus in parental (RHΔ*ku80*), knock-in and knockout parasites, 5 µg of gDNA from each line was digested with the restriction enzymes indicated in **Fig 4A-B** and **Fig S2** before hybridization with probes for the 5’UTR, CDS, or 3’UTR regions.

For the APR4-related probes, templates for probe synthesis were generated from the pTKO2_II-mE-APR4 knock-in plasmid by restriction digestions followed by gel purification. The template for the 5’UTR probe was released by *Not*I and *Avr*II digestion. The template for the CDS probe was released by *Mre*I *and Rsr*II digestion. The template for the 3’UTR probe was released by *SgrD*I and *PspOM*I digestion. After gel purification, probes were synthesized from the templates by nick translation in the presence of all four dNTPs plus biotin-dATP.

For the APR9-related probes, the following templates for probe synthesis were generated from the pTKO2_II-mE-APR9 knock-in plasmid by restriction digestions: the template for the 5’UTR probe was released by *Mfe*I and *EcoR*V digestion; the template for CDS probe 1 was released by *Sma*I and *Ngo*MIV digestion; the template for the 3’UTR probe was released by *Bstz*17I and *Nhe*I digestion. CDS probe 2 was amplified from the pTKO2_II-mE-APR9 knock-in plasmid by PCR using the primer pair S17/AS17.

### Sample preparation and imaging for Expansion Microscopy (ExM)

Expanded samples of intracellular and extracellular parasites were prepared, labeled with anti-tubulin and anti-GFP antibodies, and imaged with a DeltaVision OMX Flex imaging station (GE Healthcare-Applied Precision) as previously described [26, 30, 53].

For assessing conoid extension, freshly egressed extracellular parasites were treated with 5 µM A23187 for 1 minute at room temperature while settling on a poly-L-lysine-coated coverslip. Samples were then fixed and processed for expansion microscopy as previously described [26, 30, 53].

Sum or maximum projections were presented in the figures with contrast levels adjusted for optimization of display. The expansion ratio of ∼ 5.4 was estimated based on [30].

### Plaque assay

Freshly harvested parasites (100 or 200 per well) were used to infect confluent HFF monolayers in 6-well plates. After incubation at 37°C for seven or twelve days, the cultures were rinsed, fixed, stained, and scanned as described in [19]. Three to seven independent experiments were performed for each parasite line.

### Invasion assays

Immunofluorescence-based invasion assays were carried out as described in [19, 26] with five biological replicates. For each strain, parasites in 10 fields were counted per biological replicate. P-values for "%WT" were calculated in KaleidaGraph using two-tailed unpaired Student’s *t*-tests with unequal variance.

### Replication assay

Intracellular replication assays were performed as described in [14], with replication rates calculated as described in [54]. Three or six independent experiments were performed for each parasite line. For each strain and time point, parasites in ∼ 100 vacuoles were counted for each replicate.

### A23187-induced egress

Calcium ionophore-induced egress assays were performed as described in [19] using 5 µM A23187 in L15 imaging medium [Leibovitz’s L-15 (21083-027, Gibco- Life Technologies, Grand Island, NY) supplemented with 1% (vol/vol) cosmic calf serum].

### Electron microscopy

Suspensions of extracellular parasites were treated with A23187 and processed for negative staining as described in [19]. Samples were imaged on a Talos transmission electron microscope (Thermo Fisher) operated at 120 keV.

### Microneme immunofluorescence and secretion assays

Immunofluorescence labeling of micronemes in intracellular parasites was performed as previously described [19]. For microneme secretion assays, freshly egressed parasites were collected and treated with 5 µM A23187 as described in [55] in DMEM growth medium or L15 imaging medium. Secreted and pellet fractions were analyzed by Western blot as described in [55].

## Acknowledgments

We thank Matea Susac for tissue culture support and helpful discussions and the EM facility at Arizona State University for instrumentation support.

## Supplementary materials

**Video S1:** Time-lapse microscopy of RH*Δku80* parental (*WT*), *ΔkinesinAΔapr9* (*kinAapr9DKO*), and *ΔkinesinAΔapr9*:*mE-APR9* complement (*DKO:mE-APR9* comp) parasites. The cell impermeant DNA binding dye DAPI was added to the culture medium prior to the start of the experiment. The parasite-dependent host cell-permeabilization was detected by the DNA labeling by DAPI entering the host cell nucleus. Time is shown in min:s, Video speed: 12 frames/s. Scale bar: 5 µm.

**Fig. S1.**
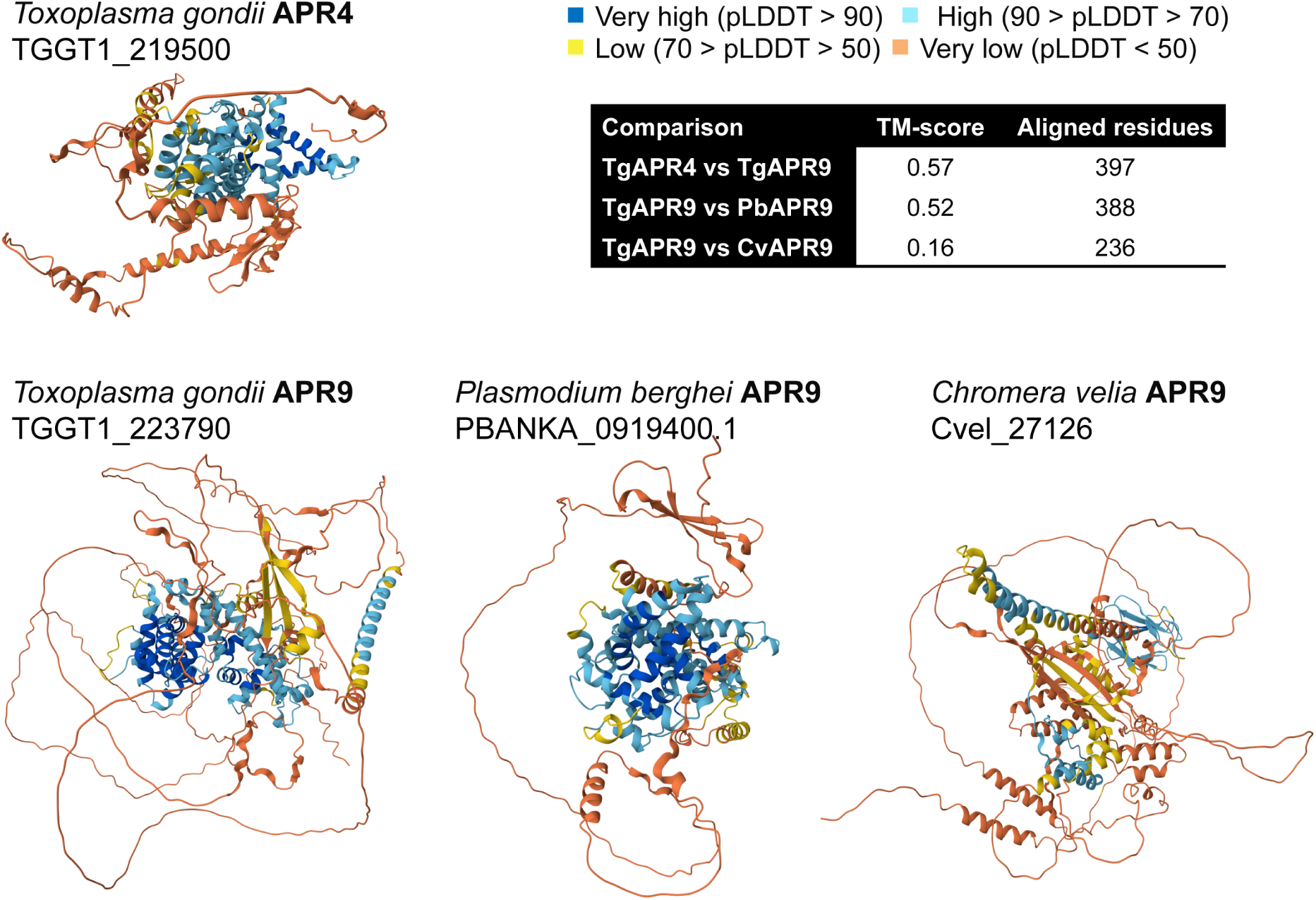
Predicted structures of TgAPR9, TgAPR4, and APR9 orthologs from *Plasmodium berghei* and *Chromera velia* by AlphaFold3. The numbers of aligned residues were generated by TM-align (Version 20190822) [57].

**Fig. S2.**
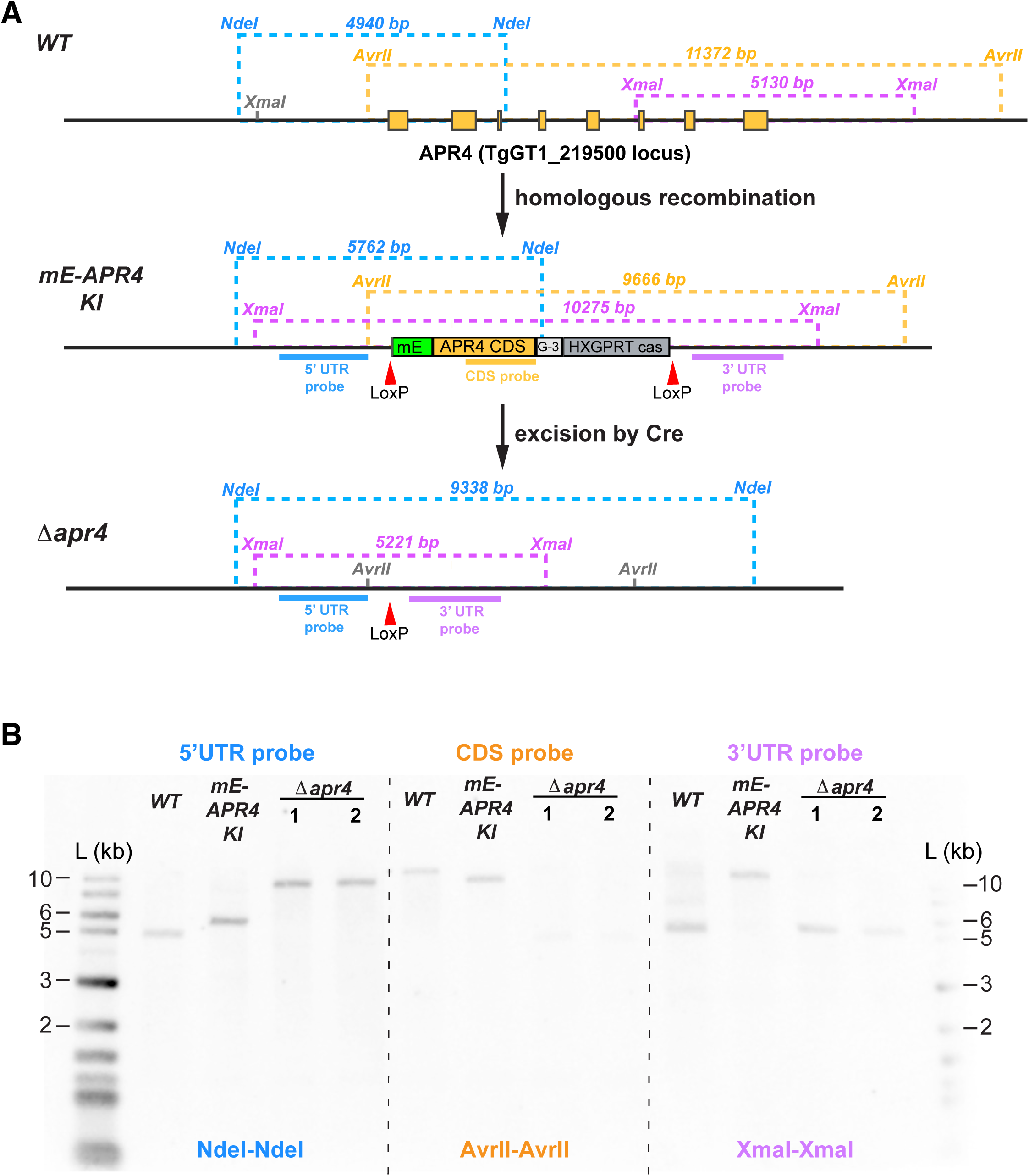
Southern blot analysis of the *apr4* locus in the WT, *mEmeraldFP-APR4* (mE-APR4 KI) and *Δapr4* parasites. **A.** Schematic for the predicted APR4 locus in WT, *mE-* tagged APR4 knock-in, and *Δapr4* lines. Restriction sites, hybridization targets of the Southern blot probes for the *apr4* coding region (CDS probe, orange bar), regions upstream (“5’ UTR probe”, blue bar) and downstream (“3’ UTR probe”, purple bar) of the CDS, and the corresponding DNA fragment sizes expected are indicated. **B.** Southern blots confirmed the homologous integration of the mE-APR4 fusion in the knock-in line, and the deletion of the *apr4* locus in the *Δapr4* lines.

**Table S1.** List of proteins identified in MudPIT analysis of an immunoprecipitation using GFP-Trap and lysate from the mEmeraldFP-APR2 knock-in line. The RH*Δku80* parental parasite having untagged APR2 was used as the negative control.

**Table S2.** Quantification of A23187-induced MIC2 secretion of WT and *ΔkinesinAΔapr9* parasites in DMEM growth medium and L15 imaging medium. Values represent background-subtracted intensities of the secreted MIC2 band in the A23187-induced samples relative to WT parasites in DMEM growth medium, normalized to the corresponding tubulin loading control in the pellet fraction.

| | Media (+A23187) | replicate 1 | replicate 2 | replicate 3 | Average ±SEM |
| --- | --- | --- | --- | --- | --- |
| RH$\Delta ku80$<br>(WT) | DMEM | 100% | 100% | 100% | 100% |
| | L15 | 172% | 224% | 77% | 158±43% |
| $\Delta kinesinA$<br>$\Delta apr9$ | DMEM | 12% | 18% | 12% | 14±2% |
| | L15 | 22% | 39% | 23% | 28±6% |

**Table S3.** List of primers used in this study.

**Table S4.** Combined raw data

