## Supplementary material for "The human parasite, *Toxoplasma gondii,* is paralyzed without two components of the apical polar ring": TableS2

**Table S2**. Quantification of A23187-induced MIC2 secretion of WT and *ΔkinesinAΔapr9* parasites in DMEM growth medium and L15 imaging medium. Values represent background-subtracted intensities of the secreted MIC2 band in the A23187-induced samples relative to WT parasites in the DMEM growth medium, normalized to the corresponding tubulin loading control in the pellet fraction.

| | Media (+A23187) | replicate 1 | replicate 2 | replicate 3 | Average ±SEM |
|---|---|---|---|---|---|
| *RHΔku80* (WT) | DMEM | 100% | 100% | 100% | 100% |
| | L15 | 172% | 224% | 77% | 158±43% |
| *ΔkinesinA Δapr9* | DMEM | 12% | 18% | 12% | 14±2% |
| | L15 | 22% | 39% | 23% | 28±6% |
